# CENP-F couples cargo to growing and shortening microtubule ends

**DOI:** 10.1101/083618

**Authors:** Gil Kanfer, Martin Peterka, Vladimir K. Arzhanik, Alexei L. Drobyshev, Fazly I. Ataullakhanov, Vladimir A. Volkov, Benoît Kornmann

**Author notes:** these authors contributed equally (Gil Kanfer, Martin Peterka, Vladimir K. Arzhanik). these authors contributed equally (Vladimir A. Volkov, Benoît Kornmann).

## Abstract

Dynamic microtubule ends exert pulling and pushing forces on intracellular membranes and organelles. However, the mechanical linkage of microtubule tips to their cargoes is poorly understood. CENP-F is a non-motor microtubule-binding protein that participates in microtubule binding at kinetochores and in the mitotic redistribution of the mitochondrial network. CENP-F-driven mitochondrial transport is linked to growing microtubule tips, but the underlying molecular mechanisms are unknown. Here we show that CENP-F tracks growing microtubule ends in living cells. In vitro reconstitution demonstrates that microtubule tips can transport mitochondria and CENP-F-coated artificial cargoes over micrometer-long distances, during both growing and shrinking phases. Based on these and previous observations, we suggest that CENP-F might act as a transporter of mitochondria and other cellular cargoes by attaching them to dynamic microtubule ends.

## Introduction

Mitochondria form a highly dynamic network. Their intracellular transport and distribution is achieved through interaction with the cytoskeleton (Okamoto & Shaw, 2005). In animal cells, microtubules are the main tracks for mitochondrial transport. The classical model for organelle transport states that adaptor proteins on the surface of the organelle recruit molecular motors, such as kinesins and dynein to transport mitochondria along the lattice of microtubules. In the case of mitochondria, the outer-membrane protein Miro binds to both the kinesin KIF5b and the dynein-dynactin complex through the cytosolic adaptor Milton (Trak1 and Trak2), thus fulfilling the requirement for mitochondrial transport (Birsa *et al*, 2013; Stowers *et al*, 2002; Guo *et al*, 2005). Recently, we have discovered that at the end of mitosis, Miro also binds to the Centromeric protein F (CENP-F), thereby influencing mitochondrial distribution in daughter cells (Kanfer *et al*, 2015). CENP-F is a non-motor microtubule-binding protein which is expressed with strict cell-cycle dependency (Zhu *et al*, 1995; Feng *et al*, 2006). Absent in G1, it accumulates in the nucleus and at the mitochondria during the S and G2 phases. During early mitosis (prophase), CENP-F localizes to the nuclear envelope and participates in its disassembly (Bolhy *et al*, 2011). In prometaphase and until anaphase, CENP-F localizes to the outer kinetochore and participates in linking kinetochores to spindle microtubules (Liao *et al*, 1995). During late mitosis and early G1, CENP-F is recruited by Miro to the mitochondrial network to participate in its proper distribution (Kanfer *et al*, 2015).

Unlike classical mitochondrial movement, which happens along the microtubule lattice, CENP-F/Miro-driven movements are linked to growing microtubule tips. Coordinated events of microtubule growth and mitochondrial movements are observed in wild type cells, but disappear in CENP-F-deficient cells. These observations together suggest that, in addition to using microtubules as passive tracks for motor-based transport, mitochondrial distribution in mitosis is linked to microtubule dynamics.

Both growth and shrinkage of microtubules produce mechanical force. The force generated by growing microtubules can be observed *in vitro* when microtubules are grown toward a wall (Kerssemakers *et al*, 2006). These forces can also be harnessed by one factor, the tubulin polymerase XMAP215 to move a microbead in a reconstituted *in vitro* system (Trushko *et al*, 2013). The force generated by microtubule shrinking can be harnessed by a number of proteins *in vitro* (Lombillo *et al*, 1995; Asbury *et al*, 2006; McIntosh *et al*, 2008; Welburn *et al*, 2009; Grissom *et al*, 2009). Most of these proteins were found on the kinetochores, suggesting that microtubule depolymerization is a determinant of genome segregation. Among these, CENP-F has been recently shown to harness microtubule depolymerization and transmit piconewton amounts of mechanical force to move a microbead *in vitro* (Volkov *et al*, 2015). This raises the question whether CENP-F-dependent mitochondrial movements can be directly driven by microtubule dynamics, without involving molecular motors.

## Results

### CENP-F tracks growing microtubule tips *in vivo*

We sought to image CENP-F-based mitochondrial movements by time-lapse microscopy. However, ectopic expression of N- or C-terminal GFP fusions of CENP-F did not recapitulate its endogenous localization. This was possibly due to interference of GFP with CENP-F’s microtubule-binding domains, which are located on both termini of the protein. To circumvent this, we used an internal tagging strategy to insert GFP after amino acid 1529 in between two coiled-coiled regions of CENP-F (Fig. 1A). Ectopic expression of this internally tagged CENP-F construct (hereafter referred to as CENP-F^^^GFP) in live cells faithfully recapitulated the localization of endogenous CENP-F, as it localized to mitochondria, nucleus, nuclear envelope and kinetochores, depending on the cell cycle as expected (Fig. 1B-C). We have previously reported by immunofluorescence in fixed cells that CENP-F localizes to EB1-labelled microtubule tips (Kanfer *et al*, 2015). This localization pattern was however not readily visible in live cells, probably because it was masked by free CENP-F^^^GFP. We therefore turned to Total Internal Reflection Fluorescence (TIRF) microscopy to image only microtubules in proximity to the coverslip. In these conditions, we observed many CENP-F^^^GFP foci localizing to growing microtubules (Fig. 1D). Moreover, these foci moved together with EB1 comets in the direction of of microtubule growth (Fig. 1E, Video 1), indicating that CENP-F tracks growing microtubules. Like endogenous CENP-F (Kanfer *et al*, 2015), CENP-F^^^GFP localized ahead of the EB1 comets, indicating a localization at the extreme tips of microtubules. CENP-F tracked microtubules for shorter distances and shorter times on average than EB1, but the speed of CENP-F-positive comets was no different from CENP-F-negative EB1-positive comets (Fig. 1F).

**Figure 1.**
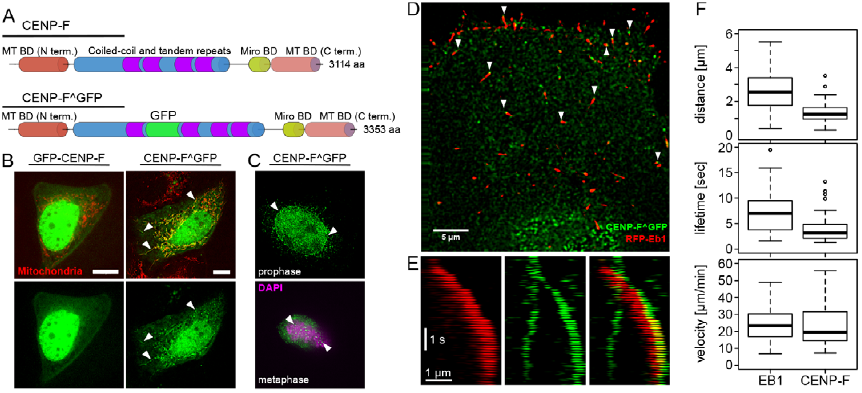
CENP-F tracks growing microtubules *in vivo.* (A) Top, domain organization of CENP-F. Bottom, position of the internal GFP-tag (CENP-F^GFP). (B) Live images U2OS cells co-expressing N-terminally GFP-tagged CENP-F (GFP-CENP-F, left) or internally GFP-tagged CENP-F (CENP-F^GFP, right), and a mitochondrial marker (mtBFP, red). Arrowheads show mitochondrial localization. Scale bar: 10 µm (C) Top, localization of Cenp-f^GFP to the nuclear envelope (arrowheads) in early prophase. DNA is counterstained with DAPI. Bottom, localization of CENP-F^GFP to the kinetochores (arrowheads). Condensed chromatin is counterstained using an anti phospho-histone H3 (H3P). Scale bars: 10 µm (D) TIRF images of live U2OS cells co-expressing CENP-F^GFP (green) and RFP-EB1 (red). Arrowheads point at CENP-F foci found at extreme tips of growing microtubules. Scale bar: 5 µm. (E) Kymograph of the live images in (D). Scale bars: 1 µm and 1 s. (F) Quantification of the distance travelled (top), lifetime (middle) and speed of comets labeled with EB1 (left) and CENP-F (right).

Although we were able to visualize CENP-F^^^GFP on mitochondria and on growing microtubule tips, we were however not able to record CENP-F^GFP moving with growing microtubule tips attached to mitochondria. One possible explanation is that, while the use of TIRF microscopy is necessary to visualize microtubule-bound CENP-F, CENP-F-driven mitochondria-microtubule interactions might happen outside of the evanescent field of total internal reflection. This effect might be aggravated by the cell-cycle regulation CENP-F/Miro interaction. Indeed, CENP-F is most present at the mitochondria during late mitosis (Kanfer *et al*, 2015), a phase when cells adopt a rounded non-adherent morphology.

### CENP-F can couple cargo transport to microtubule dynamics *in vitro*

Because of these technical limitations, we turned to a reconstituted *in vitro* system. We prepared crude mitochondria from U2OS cells overexpressing Miro1 and a mitochondria-targeted fluorescent marker (mtBFP). These mitochondria were incubated with a previously characterized recombinant fragment of CENP-F (Volkov *et al*, 2015) encompassing the Miro-binding domain (Kanfer *et al*, 2015), and added to dynamic microtubules. Dynamic microtubule extensions were grown on coverslips using fluorescently labeled GTP-tubulin and microtubule seeds, which were stabilized by the GTP analog GMPPCP (which is essentially non hydrolyzable within the time course of our experiments), and anchored to the coverslip via anti-digoxigenin IgG (Fig. 2B). The recombinant CENP-F fragment, sfGFP-2592, contains the C-terminal amino acids 2592-3114 of CENP-F, harboring both microtubule and Miro-binding domains, N-terminally fused to a superfolder GFP (sfGFP) (Pédelacq *et al*, 2006). Using TIRF microscopy, we observed that mitochondria harboring both blue (mtBFP) and green (CENP-F-sfGFP-2592C) fluorescence bound to the microtubule extensions, and moved with the tips of both growing and shortening microtubules (Fig 2, Video 2). Mitochondrial particles moved with shrinking microtubules over 2.4 ± 0.4 µm (n = 16, mean ± SEM) and with growing microtubules over 0.8 ± 0.1 µm (n = 11). Thus microtubule-dynamics-linked mitochondrial transport can be reconstituted in an *in vitro* setup, suggesting that CENP-F can mediate mitochondrial transport by direct attachment to growing and shrinking microtubule tips.

**Figure 2.**
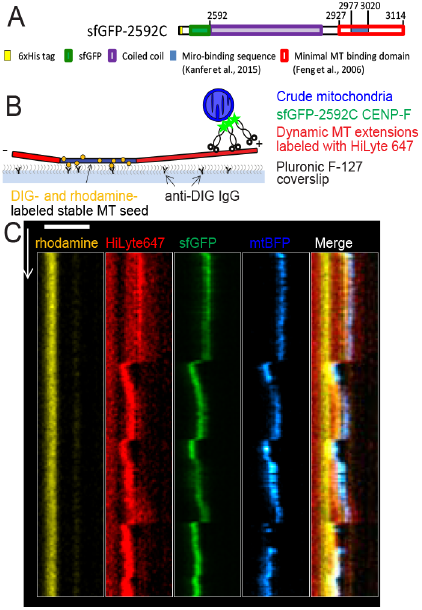
*In vitro* reconstitution of mitochondria MT-tracking using CENP-F fragment. (A) Domain organization in recombinant sfGFP-2592C fragment of human CENP-F. Numbers show aminoacid position in the full-length human CENP-F. (B) Schematics of the experimental setup. GMPCPP-stabilized, rhodamine- and DIG-labeled microtubule seeds are immobilized on a coverslip by anti-DIG antibodies. Dynamic microtubules are extended from these seeds by incubation with HiLyte-647-labeled tubulin and GTP. Crude mitochondria expressing a fluorescent marker (mtBFP) are incubated with recombinant sfGFP-2592C and added to the flow chamber. (C) Kymograph showing microtubule seed (orange), dynamic microtubule extension (red), sfGFP-2592C (green) and a mitochondrial particle (blue). Horizontal scale bar: 2 µm, vertical scale bar: 60 s.

However, to exclude the potential influence of other factors that might co-purify with mitochondria, such as kinesins and dynein, we turned to a cleaner system where mitochondria were mimicked by CENP-F-coated 1 µm glass microbeads (Fig. 3A). We used a laser trap to bring beads coated with non-fluorescent 2592C fragment into the vicinity of the tips of dynamic microtubules, released the beads and followed beads and microtubule dynamics using differential interference contrast (DIC) optics (Fig. 3B). Using this assay, we could record beads following both microtubule growth and shrinkage (Video 3). 60 out of 209 beads coated with 2592C successfully bound to microtubules (Table 1). Among the bound beads, 56% did not show any movement, 12% diffused along the microtubule lattice and 32% moved with microtubule dynamics; 18% followed only microtubule shortening, 2% (1 bead) followed only microtubule growth and 12% stayed attached to the microtubule tip during both growth and shortening through a total of 12 catastrophe events. The ability to follow microtubule tip dynamics over multiple cycles of growth and shortening was also observed for beads coated with GFP-tagged version of the C-terminal fragment (sfGFP-2592C), indicating that the sfGFP tag did not impede movement with microtubule growth (Fig 3C, Table 1 and Video 4).

**Figure 3.**
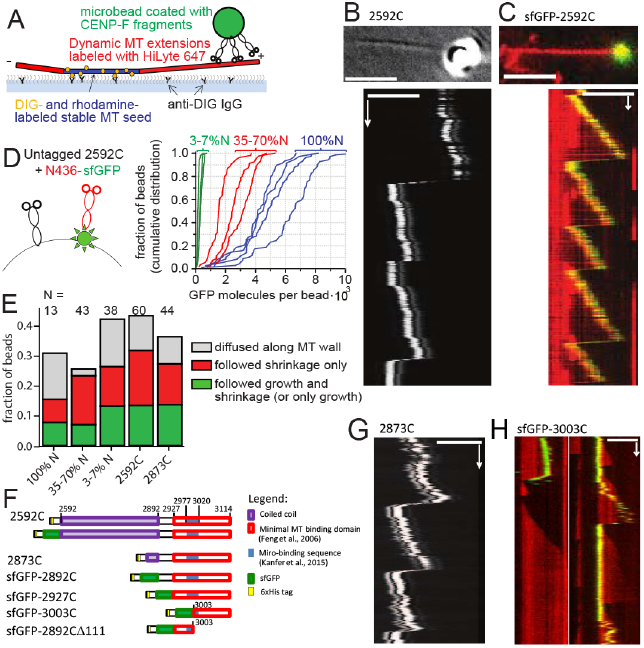
Transport of CENP-F coated beads by dynamic MT tips in vitro. (A) Experimental setup is analogous to Figure 3A, except that CENP-F coated beads are used instead of mitochondria. (B) Top: DIC image showing a 1 µm glass bead coated with 2592C on a growing microtubule tip (averaged from 20 sequential frames). Bottom: kymograph showing movement of the same bead with the microtubule tip. (C) Top: 1 µm glass bead coated with sfGFP-2592C (green) on a dynamic microtubule labeled with HiLyte-647 (red). Bottom: kymograph showing movement of the same bead. (D) Mixtures of untagged 2592C and sfGFP-tagged N436 were added to the beads (left, not to scale). The graph shows cumulative distributions of individual bead fluorescence for three molar ratios of 2592C and N436-sfGFP: 1:10 (green) 1:1 (red) and only N436-sfGFP (blue). Intensity of a single GFP molecule was determined from the bleaching curves of individual N436-sfGFP molecules as described in Materials and Methods. (E) Fate of the beads that were successfully bound to dynamic microtubules depending on the bead coating. (F) Domain organization of the C-terminal fragments of CENP-F used. Numbers show aminoacid position in the full-length human CENP-F. (G) Kymograph showing the movement of a 1 µm glass bead coated with 2873C and attached to the tip of dynamic microtubule by means of an optical trap; the bead and microtubule dynamics were imaged using DIC optics. (H) Beads coated with sfGFP-3003C (green) were allowed to bind spontaneously to dynamic microtubules grown using HiLyte-627-labeled tubulin (red). Images show two representative kymographs. Scale bars: 60s (vertical); 5 µm (horizontal).

**Table 1.** Quantification of binding and motility of beads coated with CENP-F fragments.

| Construct: | 2592C (laser trap assay) | 2873C (laser trap assay) | sfGFP-2592C | sfGFP-2892C | sfGFP-2927C | sfGFP-3003C | sfGFP-2892CΔ111 |
| --- | --- | --- | --- | --- | --- | --- | --- |
| Moved with growth | 8 | 6 | 5 | 12 | 14 | 7 | 0 |
| Moved with shortening | 18 | 12 | 21 | 34 | 14 | 9 | 2 |
| Total bound beads | 60 | 44 | 36 | 60 | 29 | 27 | 10 |
| Total beads | 209 | 164 | 97 | 358 | 300 | 567 | 424 |
| Fraction of beads binding | 0.29 | 0.27 | 0.37 | 0.17 | 0.1 | 0.05 | 0.02 |

Human CENP-F contains two distinct microtubule-binding sites that bind preferentially to different tubulin oligomers (Volkov *et al*, 2015). To mimic a possible complementing action of N- and C-terminal microtubule-binding sites of the whole length CENP-F we added various mixtures of non-fluorescent 2592C (“C”) and N436-sfGFP (“N”) to the beads, resulting in the dilution of GFP signal on the beads as a function of C:N ratio (Fig. 3D). No synergistic improvement of microtubule binding was observed at any concentration. Beads coated with 35-100% N436-sfGFP moved with microtubule growth in 7% of cases, and addition of more 2592C increased this fraction to 13% (Fig. 3E). Thus the C-terminal 2592C CENP-F fragment alone was sufficient to provide tip-tracking ability to microbeads.

We then wondered what structural features in the C-terminal fragment of CENP-F were responsible for tip tracking. To address this point we made a series of truncations in the construct and assessed their ability to attach beads to MT tips. The first feature we addressed was the long coiled-coil domain present in the 2592C fragment. Long coil-coiled domains have been shown to play important roles by positioning the cargo optimally for force transmission (Volkov *et al*, 2013). In the case of CENP-F, the long coiled-coil might mediate end-on attachment of the beads, allowing growing microtubule tip to exert a pushing force on the cargo. This conformation would however be incompatible with mitochondrial transport since the Miro-binding domain of CENP-F is situated beyond the coiled-coil domain, proximal to the microtubule-binding domain. Beads coated with a fragment of CENP-F lacking most of the coiled-coil region (2873C, Fig. 3F) could still move with microtubule growth (Fig. 3G), and their behavior was indistinguishable from the beads coated with 2592C (Fig. 3E). Therefore, coiled-coil-mediated end-on attachment of the cargo is unlikely. Consistent with this idea, the forces generated during bead movement were too small to be detected by our setup (<0.2 pN, n = 67 beads, 232 attempts). This low force is incompatible with an end-on attachment that allows microtubules to develop 2-4 pN of pushing force (Kerssemakers *et al*, 2006). We therefore conclude that CENP-F coated microbeads are more likely to be dragged behind the growing microtubule tip than pushed in front of it.

We generated two more fragments of CENP-F, one containing the previously-reported microtubule binding domain (2927-3114), and a shorter one that did not encompass the Miro-binding domain (3003-3114). Both fragments were able to promote cargo movement with growing and shrinking microtubules, although they bound microtubules more rarely than longer fragments (Table 1). As a control, a fragment missing the terminal 111 amino acids of CENP-F (2892C∆111) induced binding of only 2% of beads to the microtubules, and we did not observe any beads moving with microtubule growth (Table 1). Therefore, microtubule-tracking properties are intrinsic to the CENP-F C-terminal microtubule binding site; neither coiled-coil domains nor the Miro-binding sequence are necessary for motility.

### CENP-F binds tightly to the growing microtubule tip

Many microtubule plus-tip tracking proteins do so by association with GTP-tubulin-binding EB proteins (reviewed in Akhmanova and Steinmetz, 2015). CENP-F, however appeared to track microtubules through a different mechanism since, 1) it did not colocalize with the EB1 comet *in vivo* but appeared to bind microtubules closer to their tips, and 2) CENP-F-coated beads followed shrinking microtubules after the loss of the GTP-tubulin cap. This prompted us to test if the microtubules-binding domains of CENP-F had tip-tracking capability in isolation. In order to assess whether single molecules or oligomers of CENP-F could follow microtubule growth and/or shrinkage, we used *in vitro* reconstitution and TIRF microscopy of recombinant sfGFP-2592C fragments (Fig. 4A).

**Figure 4.**
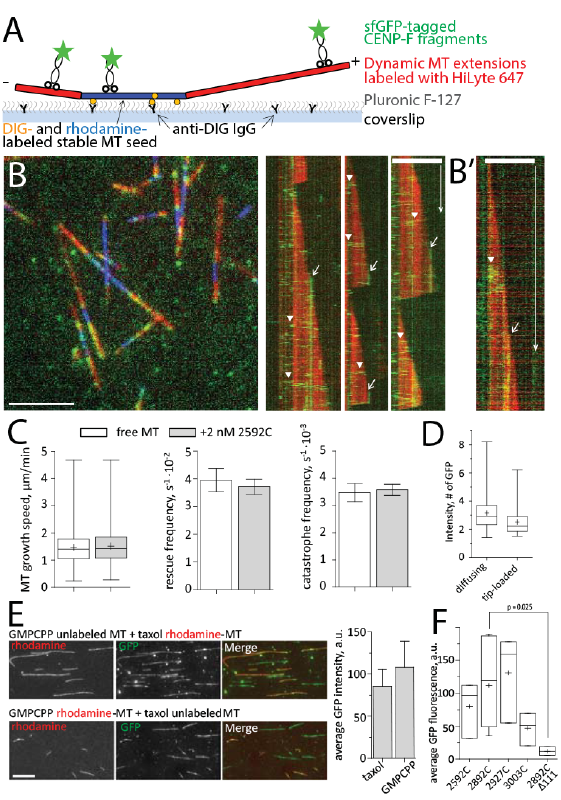
Interaction of CENP-F with dynamic MTs in vitro. (A) Schematics of experimental setup. (B) Left: dynamic MT extensions (red) growing from stable MT seeds (blue) were imaged in TIRF in the presence of 2 nM sfGFP-2592C (green). Right: kymographs showing fast-diffusing molecules of sfGFP-2592C loaded along the MT lattice (arrowheads) and immobile spots loaded at the growing MT tips (arrows). B’ shows a diffusing lattice-loaded complex (arrowhead) and tip-loaded immobile complex (arrow) imaged with a faster acquisition rate. (C) Microtubule growth speed, rescue frequency and catastrophe frequency in the absence (white bars) and in the presence (grey bars) of 2 nM 2592C. Data were summarized from 531 growth and shrinkage events for free microtubules and 610 such events in the presence of CENP-F, obtained in 3 independent experiments. Here and elsewhere: bar graphs show mean ± SEM, box and whiskers graphs show 25-75% (box), min and max (whiskers), median (horizontal line) and mean (+). (D) Intensity of 109 immobile tip-loaded and 166 diffusing lattice-loaded molecules of sfGFP-2592C in dynamic microtubule assay. (E) Flow chambers with a mixture of unlabeled GMPCPP-stabilized and rhodamine-labeled taxol-stabilized microtubules (top row), or *vice versa* (bottom row) were incubated with sfGFP-2592C. The graph shows quantification of the microtubule-associated GFP fluorescence. The values are summarized from 432 taxol-stabilized microtubules and 1018 GMPCPP-stabilized microtubules measured in 4 independent experiments. (F) Microtubule-associated GFP intensity in the presence of truncated fragments of CENP-F C-terminus (see also Fig. 4F). Data are from 3 independent experiments for sfGFP-2592C, sfGFP-2927C and sfGFP-3003C; 11 experiments for sfGFP-2892C and 2 experiments for sfGFP-2892∆111. P-value is reported based on a Mann-Whitney-Wilcoxon Rank Sum test. Scale bars: 60s (vertical); 5 µm (horizontal).

Using dynamic microtubules, we did not observe single CENP-F molecules moving at growing microtubule tips. Instead we observed two distinct behaviors of CENP-F fragments on microtubules (Fig. 4B): (1) some GFP-tagged molecules bound the microtubule lattice and diffused rapidly along microtubules before dissociating (Fig. 4B, arrowheads); and (2) some GFP-tagged molecules loaded at the growing microtubule tip, but did not follow the growth (Fig. 4B, arrows). Instead, they remained stable at their loading site and stayed bound until microtubule shortening (Video 5). Thus, individual microtubule-binding fragments of CENP-F did not appear to have intrinsic tip-tracking capability in our *in vitro* system. Previous studies showed that CENP-F fragment influences microtubule dynamics (Moynihan *et al*, 2009; Pfaltzgraff *et al*, 2016). We therefore quantified the growth and shrinkage speed of microtubule in the presence of sfGFP-2592C. Consistent with our *in vivo* data (Fig. 1F), there were no significant changes in microtubule growth rates, nor in the frequencies of rescues and catastrophes (Fig. 4C) at nanomolar concentrations of CENP-F. Therefore, CENP-F binding at the growing microtubule tip does not affect the addition of incoming tubulin dimers, suggesting that CENP-F and tubulin are not competing for the same site on microtubules.

The observed lower mobility of tip-loaded CENP-F complexes might be a result of these complexes being oligomerized and thus forming more bonds with the microtubule than the lattice-loaded ones. To test this possibility, we compared the brightness of the unbleached GFP signal of the tip-loaded complexes with the ones that loaded on the microtubule lattice. Tip-loaded complexes were not brighter than lattice-loaded complexes (Fig. 4D), suggesting that difference in mobility was not associated with a difference in oligomer size. To test whether the nucleotide associated with tubulin played any role in the binding of CENP-F, we compared the brightness of GFP signal associated with taxol-stabilized microtubules (mimicking GDP-tubulin) to the brightness of GFP signal on the GMPCPP-stabilized microtubules (mimicking GTP-tubulin, Fig. 4E). No difference was observed suggesting that CENP-F binding the tubulin-GTP cap was unlikely to result in a tighter binding of CENP-F to the growing microtubule tips. Truncated fragments (Fig. 3F) bound to stabilized microtubules with similar affinities, except for the 2892C∆111 fragment, whose binding to microtubules was barely detectable (Fig. 4F).

Finally, we tested the stability of tip-loaded and lattice-loaded complexes in high salt concentrations. To allow CENP-F loading specifically to the MT lattice, we grew microtubules from coverslip-attached seeds in the absence of sfGFP-2592C and stabilized them by addition of a GMPCPP-tubulin cap, before washing out excess tubulin. These stable microtubules were incubated in the presence of sfGFP-2592C before being washed with 100mM KCl (Fig. 5A, “lattice only”). To achieve CENP-F loading on both MT tip and lattice, we used the same experimental setup but additionally included sfGFP-2592C during the polymerization phase (Fig. 5A, “tip and lattice”). As reported previously, 100 mM KCl removed the majority of microtubule lattice-bound complexes (Volkov *et al*, 2015) (Fig. 5 B-C, “lattice only”). However, if sfGFP-2592C was present during microtubule growth, KCl wash did not result in the dissociation of microtubule-bound complexes (Fig. 5B-C, “tip and lattice”). Thus, CENP-F oligomers loaded at the growing microtubule tip formed more stable bonds with microtubules than lattice-loaded oligomers of the same size.

**Figure 5.**
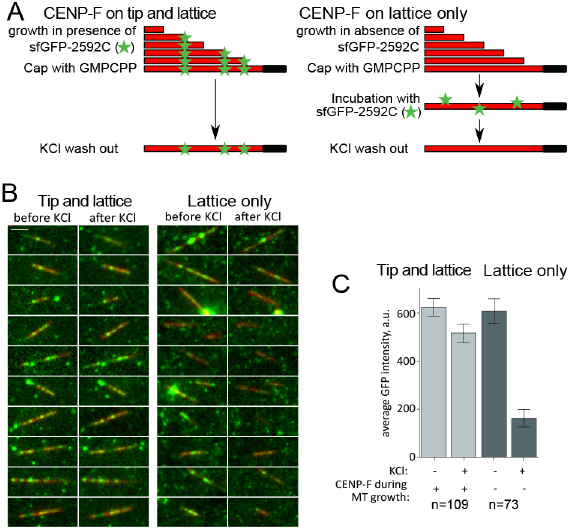
Stability of lattice- and tip-loaded molecules of sfGFP-2592C in the presence of KCl. (A) Experimental setup showing the sequence of incubations to induce a mixture of lattice- and tip-binding molecules, or only lattice-binding ones. Microtubules are grown in the presence (left) or in the absence (right) of 2 nM of sfGFP-2592C, and then stabilized by the addition of a GMPCPP-tubulin cap. Stabilized microtubules are then incubated in the same concentration of sfGFP-2592C, and washed in 100 mM KCl. The GFP signal remaining on the microtubules is then quantified. (B) Images of individual microtubules (red) grown in the presence (left) or absence of sfGFP-2592C (right) before and after KCl wash. (C) shows average microtubule-associated GFP intensity for the conditions shown in (B).

## Discussion

The spatial and temporal distributions of many cellular structures, including mitochondria, ER, secretory vesicles and chromosomes are dependent on microtubule-based transport. While chromosome movement has long been linked to microtubule dynamics, mitochondria are thought to utilize microtubules as passive tracks for kinesin- and dynein-mediated motility (Nangaku *et al*, 1994; Pilling *et al*, 2006; Birsa *et al*, 2013). Here we present direct evidence that dynamic microtubule tips can transport cargo by a motor-independent mechanism mediated by the C-terminal microtubule-binding domain of CENP-F; a glass bead coated with 2592C can follow both growing and shortening microtubule tips (Fig. 3).

Disassembling microtubules produce a power stroke due to protofilament bending, a conformational change that allows protofilaments to pull on an attached cargo (Grishchuk *et al*, 2005). But what makes a cargo move with growing microtubule tips, which are less bent, and therefore structurally much less different from the wall of the microtubule? Several mechanisms have been suggested. First, the microtubule-binding molecules might have increased affinity for the GTP cap at the growing microtubule tip, as was shown for the EB family (Zhang *et al*, 2015; Duellberg *et al*, 2016). Second, the cargo might be brought to microtubule tip by the ATP-dependent motility of a kinesin domain and track the growing microtubule tip due to the activity of an additional microtubule-binding domain, as shown for the mitotic kinesin CENP-E (Gudimchuk *et al*, 2013).

Considering that CENP-F does not localize to EB1 comets, that the C-terminal microtubule-binding site of CENP-F does not have an increased affinity for GTP-tubulin (Fig. 4E), and that molecular motors are absent in our *in vitro* system, the tip-tracking mechanism promoted by CENP-F appears to involve a third mechanism, more akin to the one that allows the microtubule polymerase XMAP215 to transport microbeads against an external load (Trushko *et al*, 2013). Indeed, the localization patterns of both CENP-F and XMAP215 at the extreme tips of microtubules are virtually indistinguishable *in vivo* (Kanfer *et al*, 2015; Nakamura *et al*, 2012), suggesting that both factors may bind similar determinants. Three major differences remain however; first, XMAP215 is not known to transport any cargo; second, CENP-F fragments alone, unlike XMAP215 (Brouhard *et al*, 2008), cannot track growing microtubule tips, despite being preferentially loaded there; third, unlike XMAP215, CENP-F binding does not influence microtubule dynamics.

The behavior of free CENP-F fragments *in vitro* is different from the behavior of full length CENP-F *in vivo.* Nonetheless, protein fragments loading onto the growing microtubule tips and staying associated there without tip-tracking is an unprecedented behavior. It suggests that CENP-F microtubule-binding domains bind structurally differently to microtubule walls and tips, and that this difference remains even after the tip becomes a wall. Since CENP-F fragments loaded at the growing microtubule tip were less mobile and less sensitive to high salt wash than the lattice-loaded ones, we suggest that terminal tubulin dimers at the growing microtubule tip expose molecular interfaces that promote higher affinity of CENP-F. Diffusion of CENP-F-coated cargo along the microtubule will then be biased towards these terminal high-affinity sites producing net movement with growing microtubule tip. The difference in behavior between free and bead-coupled CENP-F fragments suggests that CENP-F might couple organelle movement to the microtubule growth by a novel, cargo-dependent mechanism that involves collective action of multiple CENP-F molecules.

## Materials and Methods

### DNA constructs

The CENP-F^^^GFP construct was cloned by Gap-repair cloning in yeast. First, the full-length CENP-F was amplified as two separate fragments (1-1529 and 1530-3114) from a GFP-CENP-F construct (Gurden *et al*, 2010) using primers #1 and #2 for 1-1529 and #3 and #4 for 1530-3114 (Table 2). These primers introduced regions homologous with yeast vector pRS426 on both termini of CENP-F. GFP was amplified from the same vector using primers #5 and #6. These primers introduced regions homologous with CENP-F fragments generated in the previous step. The yeast episomal vector pRS426 (Sikorski & Hieter, 1989) was linearized by EcoRI digestion. For homologous recombination assembly, all four DNA fragments (CENP-F 1-1529, CENP-F 1530-3114, GFP and linearized pRS426) were co-transformed into a yeasts (BY4741) using standard procedures, which were subsequently plated on SD –URA plates to select for successful recombination events. The resulting vector carrying full-length CENP-F^^^GFP was rescued from yeast and amplified in DH5α E.Coli strain. Finally, CENP-F^^^GFP was excised using BamHI and NotI digestion and cloned into pcDNA5/frt/to mammalian expression vector, digested with the same enzymes. The final CENP-F^^^GFP construct was verified by sequencing.

Fragments of human CENP-F were generated by PCR using 2592C fragment as the template and primers #9-14 (Table 2), inserted into pET28a vector using XhoI and BamHI sites, and verified by sequencing. sfGFP was amplified by PCR using primers #7 and #8, and inserted into pET28a vector using NdeI and BamHI.

### Cell culture and transfection

U2OS cells were cultured in DMEM (Life Technologies) supplemented with 10% F.C.S. and maintained at 80% confluency. All transfections were performed using Lipofectamine 3000 (Thermo Fischer Scientific). To image EB1, cells were transfected with 0.5 µg of EB1-RFP plasmid (a gift from Tim Mitchison & Jennifer Tirnauer (Addgene plasmid # 39323)). For CENP imaging, cells were transfected with 1.5 µg of CENP-F^^^GFP plasmid. Cells were analyzed 48hrs post-transfection.

### TIRF live imaging

CENP-F^^^GFP and RFP-EB1 were imaged live at 37°C using a DeltaVision OMX 3D-SIM Super-Resolution system controlled by DV-OMX software (Applied Precision), in Ring-TIRF mode. Images were captured with an ApoN 60x / 1.4NA Oil TIRF, using 1.514 immersion oil. Images were acquired every 50-100 ms for maximum of 2 min. All experiments have been repeated a minimum of three times, with consistent results.

### Protein purification

Reagents were purchased from Sigma unless stated otherwise. Tubulin and CENP-F fragments were purified as described (Volkov *et al*, 2015). In brief, tubulin was purified from cow brains by cycles of polymerization in 0.33M Pipes and depolymerization in 50 mM MES and 1 mM CaCl2 as described (Castoldi & Popov, 2003). Cycled tubulin was then polymerized and additionally labelled with Succinimidyl Esters of 5-Carboxytetramethylrhodamine (ThermoFisher), HiLyte-647 (Anaspec) or digoxigenin (ThermoFisher) as described (Hyman *et al*, 1991). CENP-F fragments were expressed in Rosetta™ cells under induction by 100 mM IPTG for 2 hours at 37°C. Cells were lysed with ultrasound or B-PER reagent (2592C and 2873C) (ThermoFisher), and CENP-F fragments were purified on Ni-NTA-agarose beads (Qiagen), desalted and then applied on a pre-packed HiTrap column (GE Healthcare). Protein was eluted from HiTrap columns with a linear gradient from 0.1 to 1.0 M NaCl in sodium phosphate buffer at pH = 7.0 (or 7.2 for sfGFP-2892C and sfGFP-2927C). sfGFP-3003C and sfGFP-2892C∆111 were purified using single-step Ni-NTA purification. Peak fractions were determined by SDS-PAGE, aliquoted in 30% glycerol, snap-frozen in liquid nitrogen and stored at −80C.

### *In vitro* reconstitution of microtubule dynamics

Flow chambers were constructed as described previously (Volkov *et al*, 2014). Glass coverslips were cleaned in oxygen plasma (PlasmaCleaner PDC-32G) for 3 min at 400 mTorr, then immersed in a solution of 0.05% dichlorodimethylsilane in trichloroethylene for 1 hr and washed in ethanol with sonication. Microtubule seeds were assembled by incubating 75 µM unlabelled tubulin, 17 µM DIG-labeled tubulin and 1 mM GMPCPP (Jena) with or without addition of 5 µM rhodamine-labeled tubulin for 15 min at 35°C. Polymerized tubulin was sedimented at 25 000 G for 15 min and resuspended in BRB-80 (80 mM Pipes pH 6.9, 4 mM MgCl_2_ and 1 mM EGTA). After assembling the chamber using double-stick tape and silanized coverslips, the coverslip surface was incubated with BRB80 supplemented with 30 µg/ml anti-DIG IgG (Roche), then 1% Pluronic F-127, and then GMPCPP-stabilized seeds diluted 1:1600. Unbound material was washed out with 10 chamber volumes of BRB80 after each step. For laser trap experiments the motility buffer contained BRB80 supplemented with 0.4 mg/ml casein, 1 mM DTT, 12 µM unlabelled tubulin and 1.7 mM GTP. For TIRF microscopy this buffer additionally contained a glucose oxidase-catalase oxygen scavenging cascade in the presence of 10 mM DTT, and 5% of tubulin was labelled with HiLyte-647. Immediately before the start of the experiment, 15-20 µL of motility buffer were warmed up at 35°C for 25s and introduced into the chamber using a syringe or peristaltic pump. All experiments were carried out at 32°C.

### *In vitro* reconstitution of mitochondria transport by microtubules

Crude mitochondria were isolated (Wieckowski *et al*, 2009) from U2Os cells, which stably expressed mtBFP and were treated with CENP-F RNAi as described (Kanfer *et al*, 2015). The mitochondria were incubated with 50 nM sfGFP-2592C in 5 mM Hepes pH 7.4 supplemented with 250 mM sorbitol and 1 mM EGTA for 40 min at 4°C, washed twice in the same buffer by centrifugation at 10 000 G for 10 min at 4°C, and resuspended in TIRF motility buffer supplemented with 1 mM ATP. Images were acquired using a Leica DMI6000B microscope equipped with a TIRF 100x 1.47NA Oil HCX PlanApo objective and an ORCA-Flash 4.0 sCMOS camera.

**Optical trap** was custom built using an AxioImager Z1 microscope (Carl Zeiss), a Spectra Physics 5W Nd:YaG 1064 nm trapping laser, a custom-built quadrant photo detector (QPD), a 100 mW 830 nm Qioptiq iFlexx 2000 tracking laser, and a piezo-electric stage (Physik Instrumente P-561.3DD). Calibration, bead positioning and force measurement was performed using software written in-house using LabVIEW 9 as described previously (Volkov *et al*, 2013). Differential interference contrast (DIC) imaging was performed using EC Plan-Neofluar NA 1.3 objective (Zeiss) and a Cascade II EMCCD camera (Roper Scientific).

### Experiments with CENP-F-coated beads

Glass 1 µm beads coated with carboxyl groups (Bangs Laboratories) were suspended as 1% (w/v) by sonication, then activated with 50 mg/ml of Sulfo-NHS and 50 mg/ml EDC (ThermoFisher) by incubating for 30 min at 25°C in 25 mM MES pH 5.0 supplemented with 0.05% tween-20 (MES buffer). Unbound reagents were washed 3 times by sedimenting the beads for 1 min at 13 000 G. Beads were then resuspended in 3 mg/ml streptavidin (Pierce) and incubated for 2 hrs at 4°C with rocking. Reaction was stopped by addition of 20 mM glycine, the beads were washed 3 times in MES buffer and stored for several months with rocking. To attach 2592C, N436 and 2873C fragments, the streptavidin-coated beads (0.1% w/v) were functionalized with biotinylated anti-Penta His IgG (Qiagen) in PBS supplemented with 2 mM DTT, 0.4 mg/ml casein and 0.5 mg/ml BSA, washed 3 times and then incubated with the CENP-F fragment followed by 3 washes in the same buffer and resuspended in motility buffer. Fragments tagged with sfGFP were attached similarly using biotinylated anti-GFP IgG (Rockland).

For experiments with non-fluorescent 2592C and 2873C fragments, individual beads were captured in a laser trap with a spring constant of 5-10·10^−3^ pN/nm and brought in the vicinity of a growing microtubule tip. The bead was then released by shutting the trapping beam. Since our laser trap is built on an upright microscope, the beads that failed to attach sank fast out of objective focus due to the bead weight. Attached beads remained close to the coverslip and displayed arc-like motion (see Video 2). Motions of the successfully attached beads were recorded every 1-2 s. In an attempt to measure microtubule pushing force acting on the beads, the successfully attached beads were then trapped once again and bead displacement from the trap center was recorded using quadrant photo detector (QPD) until the microtubule disassembly as judged by DIC.

For TIRF microscopy of beads coated with sfGFP-tagged fragments, the flow-chambers with coverslip-attached seeds were first incubated with tubulin in motility buffer to allow microtubule extensions, and then beads resuspended in the same buffer were introduced. The fact that our TIRF microscopes were inverted allowed the beads to sink to the chamber floor with anchored microtubules and bind to microtubules spontaneously. Images were acquired using a Nikon Eclipse Ti microscope equipped with a Plan Apochromat 100x 1.49 NA objective, a TIRF laser LU-N4 module and an Andor iXon+ camera.

### Single-molecule experiments

To test whether CENP-F was tracking microtubule growth, the TIRF motility buffer was supplemented with 0.5-5 nM sfGFP-2592C. Microtubule growth and CENP-F motility were monitored using DeltaVision OMX microscope in TIRF mode equipped with ApoN 60x / 1.49NA Oil TIRF objective and four sCMOS OMX V4 cameras. Additional experiments to quantify intensity of tip-loaded and lattice-loaded complexes were performed using a Nikon Eclipse Ti TIRF microscope (see above).

### CENP-F decoration of stable microtubules

To test for nucleotide preference of CENP-F binding to microtubules, DIG-labelled, rhodamine-labelled and unlabelled tubulins were polymerized in the presence of 1 mM of GMPCPP or 10 µM taxol for 15 min at 35°C. Microtubule polymer was separated by centrifugation at 25 000G and resuspended in BRB80. Taxol stabilized rhodamine-labeled microtubules were mixed with GMPCPP-stabilized unlabelled microtubules, or taxol stabilized unlabelled microtubules were mixed with GMPCPP-stabilized rhodamine-labelled microtubules. These mixtures of microtubules were introduced into the flow chamber with silanized coverslips coated with antitubulin IgG (Serotec) and Pluronic F-127. After unbound microtubules were washed away, 2 nM of sfGFP-2592C in BRB80 supplemented with 0.4 mg/ml casein and a glucose oxidase-catalase oxygen scavenging cascade in the presence of 10 mM DTT was perfused through the chamber.

### KCl wash-out experiments

DIG-labeled GMPCPP stabilized microtubule seeds were attached to the silanized coverslips in the flow chamber as described above. The seeds were then extended by incubation with 12 µM tubulin labelled with 5% HiLyte647 with 1.7 mM GTP and with or without 2 nM sfGFP-2592C. After 10 min of microtubule growth, the dynamic tips were stabilized by rapidly exchanging the solution to the one with the same tubulin concentration, but in the presence of 1 mM GMPCPP. All tubulins were then washed out using BRB80 with 0.4 mg/ml casein and 1 mM DTT. In the control experiment that lacked sfGFP-2592C during microtubule growth, 2 nM sfGFP-2592C was added after removing tubulin. The chamber was then washed to remove unbound sfGFP-2592C and several microtubule-containing regions on the coverslip were imaged in GFP and HiLyte647 channels, and positions of these regions recorded. The chamber was then washed with the same buffer containing 100 mM KCl and the same regions on the coverslip were imaged once again.

### Image and data analysis

All microscopy images were processed using ImageJ. To correct for unevenness of the background in DIC images, at the beginning of an experiment 50-100 images in an empty chamber were collected, averaged, and a constant value of 3000 a.u. was subtracted from each pixel. The resulting average background image was subtracted from each individual DIC image acquired during this experiment. Kymographs were built using a custom ImageJ script that projects a maximum intensity across a 9-pixel-wide line. Microtubule and spot intensities were processed as described earlier (Volkov *et al*, 2014, 2015). Intensity of a single GFP molecule was obtained from the bleaching curves measured using picomolar concentrations of N436-sfGFP adsorbed to plasma-cleaned coverslips and imaged under conditions identical to the conditions in the experimental chambers. Frequencies of microtubule rescue and catastrophes were calculated as described in (Walker *et al*, 1988). Frequency of catastrophes was calculated by dividing the amount of events over total time of microtubule growth. Frequency of rescue was calculated by dividing the amount of events over total time of microtubule disassembling. Data are reported as mean ± SEM unless stated otherwise. P-values are reported based on the two-sample t-test unless stated otherwise.

## Acknowledgements

The authors are grateful to the members of Kornmann group for discussions, to Richard McIntosh (University of Colorado) for careful reading of the manuscript, to Arkady Fradkov (ZAO Evrogen, Moscow, Russia) for excellent technical assistance with cloning of CENP-F fragments. Fluorescence microscopy was performed in part at the Scientific Center for Optical and Electron Microscopy of the ETH Zurich. This study was supported by grants of the Swiss National Science Foundation (PP00P3_13365 to BK), the European Research Council (grant 337906-OrgaNet to BK), the Russian Fund for Basic Research (#15-04-04467 to FIA, #14-04-00057 to VAV), and by a fellowship from Dmitry Zimin Dynasty Foundation to VAV.

## Video Legends

**Video 1.** Time-lapse TIRF microscopy of a U2Os cell expressing RFP-EB1 (red) and Cenp-f^^^GFP (green). Arrowheads show Cenp-f^^^GFP localizing to the microtubule plus ends ahead of EB1 comets.

**Video 2.** Mitochodria isolated from U2Os cells expressing mtBFP (blue) and depleted of CENP-F using RNAi were incubated with sfGFP-2592C (green) and added to dynamic microtubules (red) growing from GMPCPP-stabilized microtubule seeds (yellow).

**Video 3.** A 1 µm glass bead coated with 2592C was attached to a growing microtubule using a laser trap. The trap was then shut down and bead and microtubule dynamics imaged using DIC optics.

**Video 4.** A 1 µm glass bead coated with sfGFP-2592C (green) spontaneously attached to a dynamic HyLite647-labeled dynamic microtubule (red).

**Video 5.** Dynamic microtubules assembled from HyLite647-labeled tubulin (red) growing from rhodamine-labeled, GMPCPP-stabilized microtubule seeds (blue) in the presence of 2 nM sfGFP-2592C (green). Arrows show tip-loading, non-motile sfGFP-2592C oligomers.

## References

Akhmanova A & Steinmetz MO (2015) Control of microtubule organization and dynamics: two ends in the limelight. Nat. Rev. Mol. Cell Biol. 16: 711–26

Asbury CL, Gestaut DR, Powers AF, Franck AD & Davis TN (2006) The Dam1 kinetochore complex harnesses microtubule dynamics to produce force and movement. Proc. Natl. Acad. Sci. U. S. A. 103: 9873–9878

Birsa N, Norkett R, Higgs N, Lopez-Domenech G & Kittler JT (2013) Mitochondrial trafficking in neurons and the role of the Miro family of GTPase proteins. Biochem. Soc. Trans. 41: 1525–1531

Bolhy S, Bouhlel I, Dultz E, Nayak T, Zuccolo M, Gatti X, Vallee R, Ellenberg J & Doye V (2011) A Nup133-dependent NPC-anchored network tethers centrosomes to the nuclear envelope in prophase. J. Cell Biol. 192: 855–71

Brouhard GJ, Stear JH, Noetzel TL, Al-Bassam J, Kinoshita K, Harrison SC, Howard J & Hyman AA (2008) XMAP215 is a processive microtubule polymerase. Cell 132: 79–88

Castoldi M & Popov A V (2003) Purification of brain tubulin through two cycles of polymerization-depolymerization in a high-molarity buffer. Protein Expr. Purif. 32: 83–8

Duellberg C, Cade NI, Holmes D & Surrey T (2016) The size of the EB cap determines instantaneous microtubule stability. Elife 5:

Feng J, Huang H & Yen TJ (2006) CENP-F is a novel microtubule-binding protein that is essential for kinetochore attachments and affects the duration of the mitotic checkpoint delay. Chromosoma 115: 320–9

Grishchuk EL, Molodtsov MI, Ataullakhanov FI & McIntosh JR (2005) Force production by disassembling microtubules. Nature 438: 384–8

Grissom PM, Fiedler T, Grishchuk EL, Nicastro D, West RR & McIntosh JR (2009) Kinesin-8 from fission yeast: a heterodimeric, plus-end-directed motor that can couple microtubule depolymerization to cargo movement. Mol. Biol. Cell 20: 963–72

Gudimchuk N, Vitre B, Kim Y, Kiyatkin A, Cleveland DW, Ataullakhanov FI & Grishchuk EL (2013) Kinetochore kinesin CENP-E is a processive bi-directional tracker of dynamic microtubule tips. Nat. Cell Biol. 15: 1079–88

Guo X, Macleod GT, Wellington A, Hu F, Panchumarthi S, Schoenfield M, Marin L, Charlton MP, Atwood HL & Zinsmaier KE (2005) The GTPase dMiro is required for axonal transport of mitochondria to Drosophila synapses. Neuron 47: 379–93

Gurden MDJ, Holland AJ, van Zon W, Tighe A, Vergnolle M a, Andres D a, Spielmann HP, Malumbres M, Wolthuis RMF, Cleveland DW & Taylor SS (2010) Cdc20 is required for the postanaphase, KEN-dependent degradation of centromere protein F. J. Cell Sci. 123: 321–30

Hyman A, Drechsel D, Kellogg D, Salser S, Sawin K, Steffen P, Wordeman L & Mitchison TJ (1991) Preparation of Modified Tubulins. Methods Enzymol. 196: 478–485

Kanfer G, Courthéoux T, Peterka M, Meier S, Soste M, Melnik A, Reis K, Aspenström P, Peter M, Picotti P & Kornmann B (2015) Mitotic redistribution of the mitochondrial network by Miro and Cenp-F. Nat. Commun. 6: 8015

Kerssemakers JWJ, Munteanu EL, Laan L, Noetzel TL, Janson ME & Dogterom M (2006) Assembly dynamics of microtubules at molecular resolution. Nature 442: 709–12

Liao H, Winkfein RJ, Mack G, Rattner JB & Yen TJ (1995) CENP-F is a protein of the nuclear matrix that assembles onto kinetochores at late G2 and is rapidly degraded after mitosis. J. Cell Biol. 130: 507–18

Lombillo VA, Stewart RJ & Richard McIntosh J (1995) Minus-end-directed motion of kinesin–coated microspheres driven by microtubule depolymerization. Nature 373: 161–164

McIntosh JR, Grishchuk EL, Morphew MK, Efremov AK, Zhudenkov K, Volkov VA, Cheeseman IM, Desai A, Mastronarde DN & Ataullakhanov FI (2008) Fibrils connect microtubule tips with kinetochores: a mechanism to couple tubulin dynamics to chromosome motion. Cell 135: 322–33

Moynihan KL, Pooley R, Miller PM, Kaverina I & Bader DM (2009) Murine CENP-F Regulates Centrosomal Microtubule Nucleation and Interacts with Hook2 at the Centrosome. 20: 4790–4803

Nakamura S, Grigoriev I, Nogi T, Hamaji T, Cassimeris L & Mimori-Kiyosue Y (2012) Dissecting the Nanoscale Distributions and Functions of Microtubule-End-Binding Proteins EB1 and ch-TOG in Interphase HeLa Cells. PLoS One 7:

Nangaku M, Sato-Yoshitake R, Okada Y, Noda Y, Takemura R, Yamazaki H & Hirokawa N (1994) KIF1B, a novel microtubule plus end-directed monomeric motor protein for transport of mitochondria. Cell 79: 1209–20

Okamoto K & Shaw JM (2005) Mitochondrial morphology and dynamics in yeast and multicellular eukaryotes. Annu. Rev. Genet. 39: 503–36

Pédelacq J-D, Cabantous S, Tran T, Terwilliger TC & Waldo GS (2006) Engineering and characterization of a superfolder green fluorescent protein. Nat. Biotechnol. 24: 79–88

Pfaltzgraff ER, Roth GM, Miller PM, Gintzig AG, Ohi R & Bader DM (2016) Loss of CENP-F Results in Distinct Microtubule Related Defects Without Chromosomal Abnormalities. Mol. Biol. Cell 27: 1990–1999

Pilling AD, Horiuchi D, Lively CM & Saxton WM (2006) Kinesin-1 and Dynein are the primary motors for fast transport of mitochondria in Drosophila motor axons. Mol. Biol. Cell 17: 2057–68

Sikorski RS & Hieter P (1989) A system of shuttle vectors and yeast host strains designed for efficient manipulation of DNA in Saccharomyces cerevisiae. Genetics 122: 19–27

Stowers RS, Megeath LJ, Górska-Andrzejak J, Meinertzhagen I a. & Schwarz TL (2002) Axonal transport of mitochondria to synapses depends on milton, a novel Drosophila protein. Neuron 36: 1063–77

Trushko A, Schäffer E & Howard J (2013) The growth speed of microtubules with XMAP215-coated beads coupled to their ends is increased by tensile force. Proc. Natl. Acad. Sci. U. S. A. 110: 14670–5

Volkov VA, Grissom PM, Arzhanik VK, Zaytsev A V., Renganathan K, McClure-Begley T, Old WM, Ahn N, Richard McIntosh J & McIntosh JR (2015) Centromere protein F includes two sites that couple efficiently to depolymerizing microtubules. J. Cell Biol. 209: 813–828

Volkov VA, Zaytsev A V. & Grishchuk EL (2014) Preparation of Segmented Microtubules to Study Motions Driven by the Disassembling Microtubule Ends. J. Vis. Exp.: e51150–e51150

Volkov VA, Zaytsev A V, Gudimchuk N, Grissom PM, Gintsburg AL, Ataullakhanov FI, McIntosh JR & Grishchuk EL (2013) Long tethers provide high-force coupling of the Dam1 ring to shortening microtubules. Proc. Natl. Acad. Sci. U. S. A. 110: 7708–13

Walker RA, O’Brien ET, Pryer NK, Soboeiro MF, Voter WA, Erickson HP & Salmon ED (1988) Dynamic instability of individual microtubules analyzed by video light microscopy: rate constants and transition frequencies. J. Cell Biol. 107: 1437–48

Welburn JPI, Grishchuk EL, Backer CB, Wilson-Kubalek EM, Yates JR & Cheeseman IM (2009) The human kinetochore Ska1 complex facilitates microtubule depolymerization-coupled motility. Dev. Cell 16: 374–85

Wieckowski MR, Giorgi C, Lebiedzinska M, Duszynski J & Pinton P (2009) Isolation of mitochondria-associated membranes and mitochondria from animal tissues and cells. Nat. Protoc. 4: 1582–90

Zhang R, Alushin GM, Brown A & Nogales E (2015) Mechanistic Origin of Microtubule Dynamic Instability and Its Modulation by EB Proteins. Cell 162: 849–59

Zhu X, Mancini M a, Chang KH, Liu CY, Chen CF, Shan B, Jones D, Yang-Feng TL & Lee WH (1995) Characterization of a novel 350-kilodalton nuclear phosphoprotein that is specifically involved in mitotic-phase progression. Mol. Cell. Biol. 15: 5017–5029

